# Modeling *ATN1* mutations in human iPSCs reveals early developmental roles of atrophin-1

**DOI:** 10.64898/2026.10.30.760413

**Authors:** Emilia Kozłowska, Natalia Ziojła, Bartosz Nowak, Paweł Joachimiak, Magdalena Dąbrowska-Maroszek, Magdalena Sikora, Oliwia Bródka, Marta Olejniczak, Agnieszka Fiszer

## Abstract

Atrophin-1 (ATN1), encoded by the *ATN1* gene, is an evolutionarily conserved transcriptional corepressor implicated in neuronal development and severe incurable disorders, including the polyglutamine (polyQ) disease - dentatorubral–pallidoluysian atrophy (DRPLA), and the neurodevelopmental syndrome CHEDDA. However, the cellular function of ATN1 and its role during early human development remain largely uncharacterized. Here, we generated an isogenic panel of human induced pluripotent stem cell (iPSC) lines, establishing a novel resource for comprehensive studies of ATN1 function and dysfunction. This panel includes a DRPLA patient-derived line (wt/mut), CRISPR-Cas9-corrected wild-type controls (wt/wt), complete *ATN1* knock-out clones (-/-), and heterozygous lines expressing either a wild-type or a mutant allele (wt/- and -/mut). While pluripotency markers remained unaffected across all variants, further characterization revealed marked proliferation deficits in *ATN1* knock-out lines and heterozygous iPSCs without wild-type allele expression. Moreover, the formation of punctate ATN1 assemblies was enhanced in iPSCs with -/mut variants. The pronounced phenotype observed in cells expressing only the mutant *ATN1* allele may result from a combination of loss-of-function and toxic gain-of-function mechanisms. Overall, the generated panel of iPSC lines provides a unique cellular platform for investigating ATN1-dependent molecular mechanisms and their contribution to neuronal development and disease.

## INTRODUCTION

Precise regulation of gene expression is crucial for establishing and maintaining cell identity during developmental patterning and in adult life. Transcriptional control relies on the integrated activity of expression modulators, including chromatin remodeling factors and transcription factors (TFs). TFs recognize specific DNA sequences and recruit particular regulatory proteins, such as coactivators and corepressors. Complexes formed by TFs and their cofactors act through promoter and enhancer DNA elements to ensure proper spatial and temporal transcriptional programs. While equable transcriptional regulation maintains cellular homeostasis, its disruption can result in developmental abnormalities or various diseases that occur in postnatal life ^1,2^.

Atrophin-1 (ATN1), encoded by the *ATN1* gene, is an evolutionarily conserved corepressor involved in transcriptional regulation, particularly within neuronal tissues during both embryonic development and adulthood ^3,4^. Nevertheless, the cellular function of ATN1 remains poorly characterized, particularly in human cells. The *ATN1* expression is markedly enriched in brain tissues, whereas in organs such as the ovary, lungs, or skeletal muscle, it is relatively low ^5–9^. It has previously been reported that ATN1 can act as a transcriptional corepressor in nuclear receptor signaling, with orphan receptors serving as TFs, including brain-specific NR2E1 (also known as TLX) ^5,10^. The ATN1-NR2E1 interaction helps maintain NSC identity and prevents premature differentiation ^11–13^. These findings support the role of ATN1 in neuronal development, but the exact molecular network through which ATN1 governs neuronal differentiation remains largely uncharacterized.

Despite their low frequency in the human population, *ATN1* mutations are linked to a spectrum of autosomal dominant neurological disorders. Missense mutations or in-frame deletions/duplications within exon 7 of the *ATN1* gene disrupt an HX motif in atrophin-1, thereby compromising protein structure and interactions to cause the neurodevelopmental disorder CHEDDA (Congenital Hypotonia, Epilepsy, Developmental Delay, Digit Abnormalities) ^14,15^. The patients with CHEDDA demonstrate developmental delay, brain and heart malformations, and epilepsy. In contrast, pathogenic expansion of CAG repeats in exon 5 of the *ATN1* gene leads to an abnormal polyglutamine (polyQ) tract in atrophin-1 and causes dentatorubral–pallidoluysian atrophy (DRPLA) ^16,17^. DRPLA belongs to the group of polyQ diseases, that include Huntington’s disease (HD) and several types of spinocerebellar ataxias (SCAs), that generally show complex molecular pathogenesis ^18,19^. In established pathogenic concept of DRPLA as a polyQ disease, mutant atrophin-1 acquires toxic properties ^20,21^. In healthy individuals, the number of CAG repeats in *ATN1* ranges from 7 to 35, with the most common variant containing 19 repeats ^22,23^. Complete penetrance for DRPLA is achieved when the number of CAG repeats exceeds 48 ^19,24^. The pathology of DRPLA shows a strong correlation between CAG repeat length and age of symptom onset ^25^. In juvenile-onset patients, diagnosed before the age of 20, symptoms such as myoclonus, seizures, ataxia, and cognitive decline are typically observed. These young patients usually carry longer CAG tracts, ranging from 70 to 90 repeats. In contrast, adult-onset DRPLA is more frequently characterized by dementia, chorea, ataxia, and psychiatric manifestations ^26,27^. Altogether, severe neurological dysfunction triggered by disruption in atrophin-1 activity highlights its crucial role in neuronal development and in maintaining neuronal function in adult cells. However, the precise molecular mechanisms, targets, and coplayers by which ATN1 regulates neuronal identity during development, health, and disease remain largely elusive due to a lack of proper *in vitro* systems to study these aspects in a human context.

To elucidate the biological functions of *ATN1* and its contribution to human neuronal development, function, and pathology, it is essential to employ appropriate *in vitro* models that capture both the genetic context and developmental dynamics. So far, human-induced pluripotent stem cells (iPSCs) have provided an invaluable model for investigating neurodevelopmental and neurodegenerative processes, as they can be differentiated into specific neuronal subtypes while maintaining the donor’s genetic background ^28,29^. In this study, we generated a novel comprehensive panel of iPSC lines suitable for ATN1 molecular network studies. Based on DRPLA patient-derived iPSCs ^30^, we established isogenic control lines harboring two wild-type alleles and complete *ATN1* loss-of-function clones. In addition, we generated iPSCs carrying either a single *ATN1* allele with a wild-type or pathogenic (expanded CAG repeats) sequence. We characterized iPSC phenotypes carrying distinct *ATN1* variants by assessing proliferation, atrophin-1 localization, and pluripotency markers. In future studies, our set of iPSC lines could serve as a versatile *in vitro* model for dissecting the molecular mechanisms underlying ATN1 function and dysfunction during neuronal differentiation.

## MATERIALS AND METHODS

### iPSC culture

The parental iPSC line (harboring 16/68 CAG repeats in the *ATN1* gene) ^30^ and all iPSC lines established in this study were maintained in StemFlex medium (Gibco, USA) supplemented with Penicillin– Streptomycin (Sigma-Aldrich, USA) on Geltrex-coated (Gibco, USA) plates at 37 °C in a humidified incubator with 5% CO_2_. The cells were passaged every 3–4 days using PBS-EDTA. All iPSC lines were routinely screened for mycoplasma contamination by PCR and confirmed to be negative.

### Genome editing and clonal selection

Genome editing of patient-derived iPSCs was performed using a CRISPR-Cas9 method, either as a ribonucleoprotein (RNP) complex for wt/wt cl.1 or *via* a plasmid encoding sgRNAs and Cas9, with or without a *piggyBac*-based selection cassette (wt/wt cl.2, -/-, and -/mut lines). Detailed experimental protocols and sgRNA target sequences are provided in the Supplementary Materials and Methods and Supplementary Table 1. Following clonal expansion, genotyping was confirmed via PCR and Sanger sequencing, while a piggyBac excision vector and fialuridine were used to remove the selection cassette from isogenic controls.

### Immunofluorescence staining

The iPSC colonies were fixed with 4% paraformaldehyde for 15 min at room temperature and washed with PBS. Fixed cells were permeabilized with 0.5% Triton X-100 (BioShop, Canada) for 5 min and subsequently blocked with 5% BSA/0.1% Tween-20 for 45 min. Cells were then incubated overnight at 4 °C with specific primary antibodies (Supplementary Table 2) diluted in blocking buffer. The following day, cells were incubated for 1 h with secondary antibodies conjugated to Alexa Fluor 488 or TRITC fluorophores. Nuclei were counterstained with DAPI (Sigma-Aldrich, USA). Coverslips were mounted using ProLong Glass Antifade Mountant (Invitrogen, USA). Images were acquired using a Zeiss Axio Observer fluorescence microscope and a Nikon Ti2 Spinning-Disk confocal microscope. Image processing and analysis were performed using ZEN Black and ZEN Lite 3.11 software (Zeiss, Germany), ImageJ 1.54, and CellProfiler 4.2.8. For ATN1 (*n* = 3) and pluripotency markers (*n* = 2) immunofluorescence staining, at least 5 images were quantified.

### Western blot

Proteins were extracted from iPSC pellets (*n =* 4) using PB buffer (60 mM Tris-base, 2% SDS, 10% sucrose, 2 mM PMSF) by heating at 95 °C twice for 5 min each. Protein concentration was determined spectrophotometrically (Denovix, USA). Equal amounts of protein (30 μg) were resolved on a NuPAGE 4–12% Bis-Tris gel (1.0–1.5 mm; Thermo Fisher Scientific, USA) at 70 V for 15 min followed by 135 V for 2.5 h at room temperature. Proteins were then transferred onto an Amersham Protran 0.4 µm nitrocellulose membrane (Cytiva, USA) using wet transfer at 30 V overnight at 4 °C. Membranes were blocked with 5% non-fat dry milk in TBS-T (0.1% Tween-20) and incubated with primary antibodies (Supplementary Table 3) diluted in 5% non-fat dry milk in TBS-T overnight at 4 °C. Subsequently, membranes were incubated for 1 h at room temperature with HRP-conjugated secondary antibodies diluted in 5% non-fat dry milk in TBS-T. Protein bands were visualized using WesternBright Quantum chemiluminescent substrate (Advansta, USA) and imaged with a G:Box system (Syngene, UK). Band intensities were quantified using ImageJ software (NIH, USA). All membranes are shown in Supplementary Files.

### Live-cell imaging and confluency analysis

Live-cell imaging of iPSC growth (*n* = 3) was performed using the IncuCyte Live-Cell Analysis System (Sartorius, Germany). Cells were seeded at a density of 6 × 10^2^ cells/cm^2^ in Geltrex-coated 96-well plates and cultured in StemFlex medium (Gibco, USA) supplemented with 10 µM Y-27632 (StemCell Technologies, Canada) for the first 24 h. Cultures were maintained in the IncuCyte system at 37 °C in a humidified atmosphere with 5% CO_2_. To quantify cell number, 30 nM SiR-DNA and 60 nM Verapamil (Tebubio, France) were added at seeding and subsequently every 2 days during medium replacement. Phase-contrast images were acquired every 2 h, and cell confluency was quantified in real time using the IncuCyte Base Analysis Software (Sartorius, Germany). Cell proliferation was evaluated by monitoring changes in cell-occupied area (% confluency) over time. The differences in the dynamics of confluency and cell number over time between the lines were quantified as the area under the curve.

## RESULTS

To investigate the role of the *ATN1* gene at the initial stages of human cell development, we generated a set of iPSC lines carrying distinct *ATN1* variants: a wild-type line, a complete *ATN1* knock-out (KO), and lines expressing either the wild-type or the mutant allele. As the parental cell line, we used a previously obtained and characterized DRPLA patient-derived iPSC line (wt/mut) carrying one normal *ATN1* allele with 16 CAG repeats, and one mutant allele with 68 CAG repeats ^30^.

### Generation of control iPSC lines

First, to correct the mutant allele, RNP complexes targeting exon 5 of the *ATN1* gene together with donor template were electroporated into iPSCs (**Fig. 1A, B, strategy 1**). We identified only a single corrected clone with retained 16 CAG at one allele and a contraction of the mutant allele from 68 to 19 CAG repeats, following Cas9-mediated cleavage (clone named as wt/wt cl.1, **Fig. 1C**). Due to the low efficiency of strategy 1, we adopted strategy 2 (**Fig. 1A, B, strategy 2**) which combined CRISPR-Cas9-mediated genome editing with a homologous recombination (HR) donor template containing a *piggyBac*-based selection cassette, as previously described ^31^. The donor construct contained homology arms of *ATN1* sequence with 15 CAG repeats, and a puromycin-resistance selection cassette. Following clonal picking, we detected a few iPSC lines containing 15 CAG repeats and a selection cassette at one allele, with 16 CAG repeats retained at the other allele. Furthermore, we removed the selection cassette sequence from one selected clone using the piggyBac excision-only vector, as confirmed by PCR with primers flanking or specific for the cassette’s sequence (**Supplementary Fig. 1A**). As a result, we obtained several clonal iPSC lines with corrected length of CAG tract in *ATN1* gene, and for further analysis, we chose one, referred here as wt/wt cl.2 (**Fig. 1C**). Western blot analysis confirmed synthesis of the full-length wild-type atrophin-1 and the absence of the mutated protein in all generated control iPSC lines (**Fig. 1D**). ATN1 level in wild-type clones was similar to the parental iPSC line, but in wt/wt cl.1 iPSCs, a tendency for increased ATN1 level was noted (*p*-value = 0.078, **Fig. 1E**).

**Figure 1.**
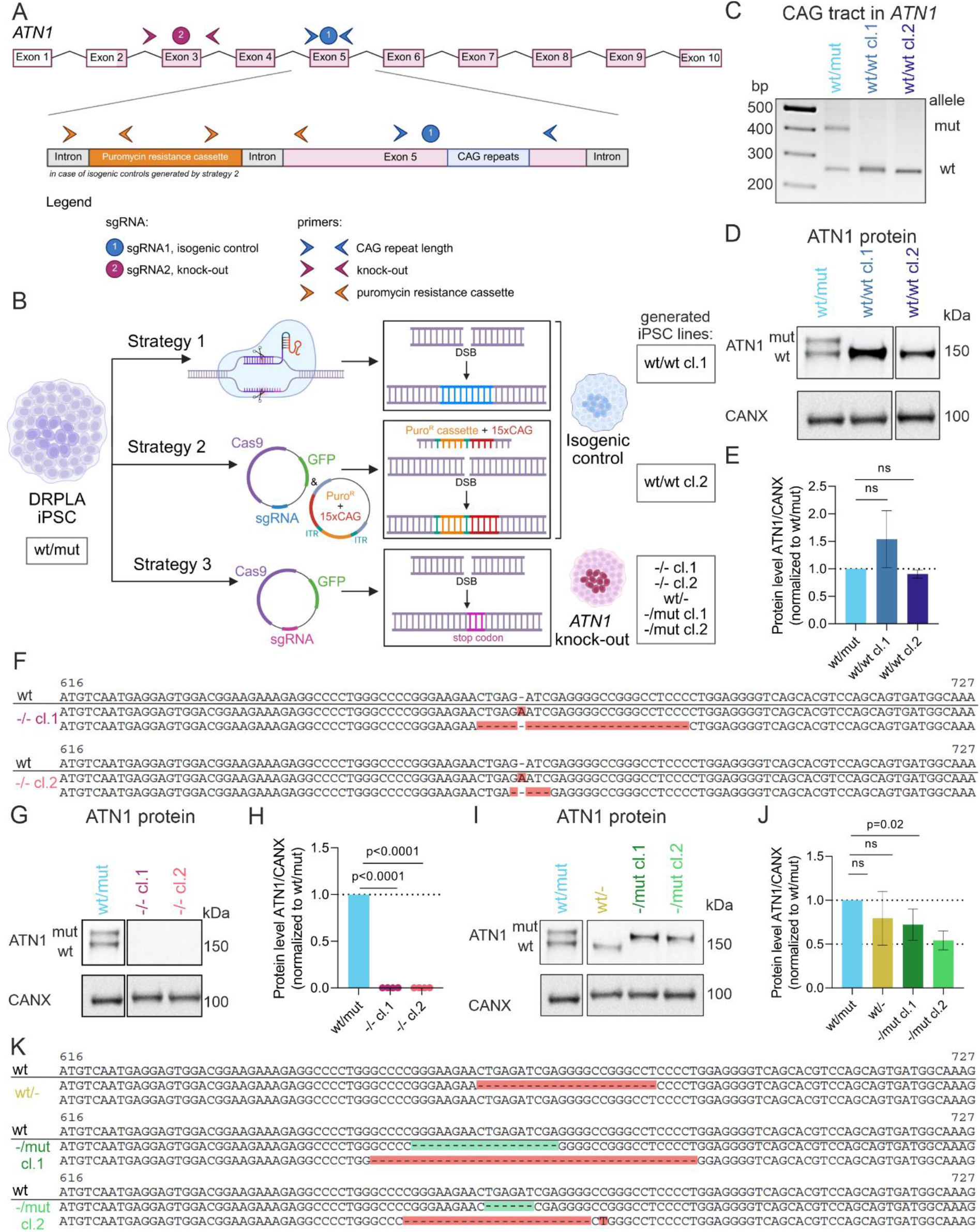
Generation of human iPSCs with different variants of the *ATN1* gene. **A**. Schematic representation of the *ATN1* gene indicating the binding sites of sgRNAs used for CRISPR-mediated editing, and primers employed for genotyping. **B**. Overview of the CRISPR/Cas9 strategies used to establish isogenic controls (wt/wt, strategy 1 and 2), complete *ATN1* knock-outs (-/-, strategy 3), and single-allele expressing iPSC lines (wt/-, -/mut, strategy 3) from parental DRPLA-patient derived iPSCs (wt/mut). **C**. PCR amplification of *ATN1* fragment spanning CAG repeat tract, in parental (wt/mut) and generated isogenic control lines (wt/wt cl.1 and wt/wt cl.2), confirming the absence of expanded tract at the genomic level. **D**. Representative western blot analysis of ATN1 protein in parental (wt/mut) and isogenic control (wt/wt cl.1 and wt/wt cl.2) iPSC lines. **E**. Quantification of western blot analysis of ATN1 level in wt/mut and wt/wt iPSCs. *n* = 4 **F**. A part of the genomic *ATN1* sequence showing the introduced indels leading to premature stop codons (red) in both *ATN1* knock-outs (-/- cl.1 and -/- cl.2) compared to the wild-type non-edited sequence (wt). **G**. Representative western blot analysis of ATN1 protein in parental (wt/mut) and *ATN1* knock-out iPSCs (-/- cl.1 and -/- cl.2). **H**. Quantification of western blot analysis of ATN1 level in wt/mut and -/- iPSCs. *n* = 4 **I**. Representative western blot analysis of ATN1 protein in parental (wt/mut) and *ATN1* single knock-out iPSCs (wt/-, -/mut cl.1 and -/mut cl.2). **J**. Quantification of western blot analysis of ATN1 level in wt/mut and *ATN1* single knock-out iPSCs (wt/-, -/mut cl.1 and -/mut cl.2). *n* = 4 **K**. A part of the genomic *ATN1* sequence showing the introduced indels leading to premature stop codons (red) or in-frame deletions (green) in iPSCs expressing only mutated or wild-type *ATN1* allele (wt/-, -/mut cl.1 and -/mut cl.2) compared to the wild-type non-edited sequence (wt). For all western blot analyses (D, G, I), full membranes are shown in the supplementary file. Calnexin (CANX) was used as a loading control. *wt* – wild-type ATN1 protein with 16 CAG, *mut* – mutated ATN1 protein with expanded 68 CAG tract. For all plots (E, H, J), data are presented as mean ± SD; for statistical analysis, one-way ANOVA for multiple comparisons (Tukey’s test) was applied, *p*-values<0.05 were considered statistically significant; *ns* – not significant. wt/mut – light blue; wt/wt cl.1 – medium blue; wt/wt cl.2 – dark blue; -/- cl.1 – dark pink; -/- cl.2 – light pink; wt/-– mustard; -/mut cl.1 – dark green; -/mut cl.2 – light green.

### Generation of *ATN1* knock-out lines

Next, to generate *ATN1* KO iPSC lines, we employed a plasmid encoding Cas9 and an sgRNA targeting exon 3 of the *ATN1* gene (**Fig. 1A, B, strategy 3**). To enhance clonal line selection, we used GFP expression from a plasmid carrying sgRNA and Cas9 to sort GFP+ electroporated cells by fluorescence-activated cell sorting (FACS), performed in bulk to improve iPSC survival. Followed by clonal picking, we identified individual iPSC clones with changes in the sequence of the *ATN1*, and for further characterization, we picked two of them, referred here as -/- cl.1 and -/- cl.2. At the DNA level, both *ATN1* knock-out clones exhibit the same mutation at one allele, specifically the 1-nucleotide (nt) insertion, resulting in frameshift and occurrence of the first premature stop codon at amino acid (AA) position 44 in exon 3. At the second allele, clones -/- cl.1 and -/- cl.2 harbor a 25-nt and 4-nt deletion, respectively, causing a premature stop codon at AA position 51 and 58 in exon 4, respectively (**Fig. 1F**). Western blot analysis confirmed the absence of ATN1 protein in both generated knock-out clones **1G, H**).

Through strategy 3, we also identified heterozygous clones expressing either the wild-type (16 CAG) or the mutant (68 CAG) allele (hereafter as wt/- and -/mut). While all wt/- and -/mut clones retained both CAG tracts at the DNA level (**Supplementary Fig. 1B**), protein synthesis was restricted to a single allele (**Fig. 1I, J**). Specifically, in the wt/-iPSC line, we observed a 22-nt deletion that introduces a premature stop codon in exon 4 at the 52^nd^ AA position at the *ATN1* allele with 68 CAG repeats. In contrast, -/mut cl.1 shows a 40-nt deletion at the allele with 16 CAG repeats leading to a premature stop codon in exon 4 at the 46^th^ AA position. In -/mut cl.2 iPSCs, we observed a 24-nt deletion and a 1-nt insertion at the wild-type allele, causing a frameshift and a premature stop codon in exon 3 at the 36^th^ AA position (**Fig. 1K**).

### Distinct ATN1 puncta frequency is increased in mutant variants

As expanded polyQ tracts often alter normal protein hallmarks, leading to misfolding, disrupted nuclear transport, or accumulation ^32–34^, we investigated whether this phenomenon is already observable in iPSCs for atrophin-1. Moreover, sequence analysis revealed that at the allele with 68 CAG repeats in -/mut clones there are additional mutations - in frame deletions leading to lack of several AA within the nuclear localization sequence (NLS) (**Fig. 2A**). Therefore, to dissect how CAG tract mutation and alterations in the NLS affect atrophin-1 protein localization in iPSCs, we performed immunofluorescence staining (**Fig. 2B**), after confirming staining specificity (**Supplementary Fig. 2A, B**). We observed predominant nuclear localization of both wild-type and mutant ATN1 as the majority of ATN1 signals, more than 60%, were from the nuclear compartment, in all investigated iPSC lines (**Fig. 2C**). When we compared ATN1 mean intensity between the wt/mut, wt/wt, and -/mut iPSC lines, we noted distinct alterations among the studied cells that the mutations may drive. Within the nucleus, the wt/mut exhibited a decreased ATN1 signal compared to wt/wt, while in the cytoplasm, the ATN1 signal was similar between these two lines. In contrast, both -/mut lines displayed a markedly elevated ATN1 signal in the cytoplasm and nucleus, relative to wt/wt line (**Fig. 2D**). These findings may suggest that the appearance of one wild-type allele may compensate for the dysfunction of the mutated allele. Subsequent precise single-cell analysis of ATN1 puncta revealed significant alterations, with -/mut clones demonstrating a significant increase in puncta frequency within both compartments compared to the control wt/wt line (**Fig. 2E**). This could result from impaired processing/interactions of the mutant atrophin-1 in -/mut clones.

**Figure 2.**
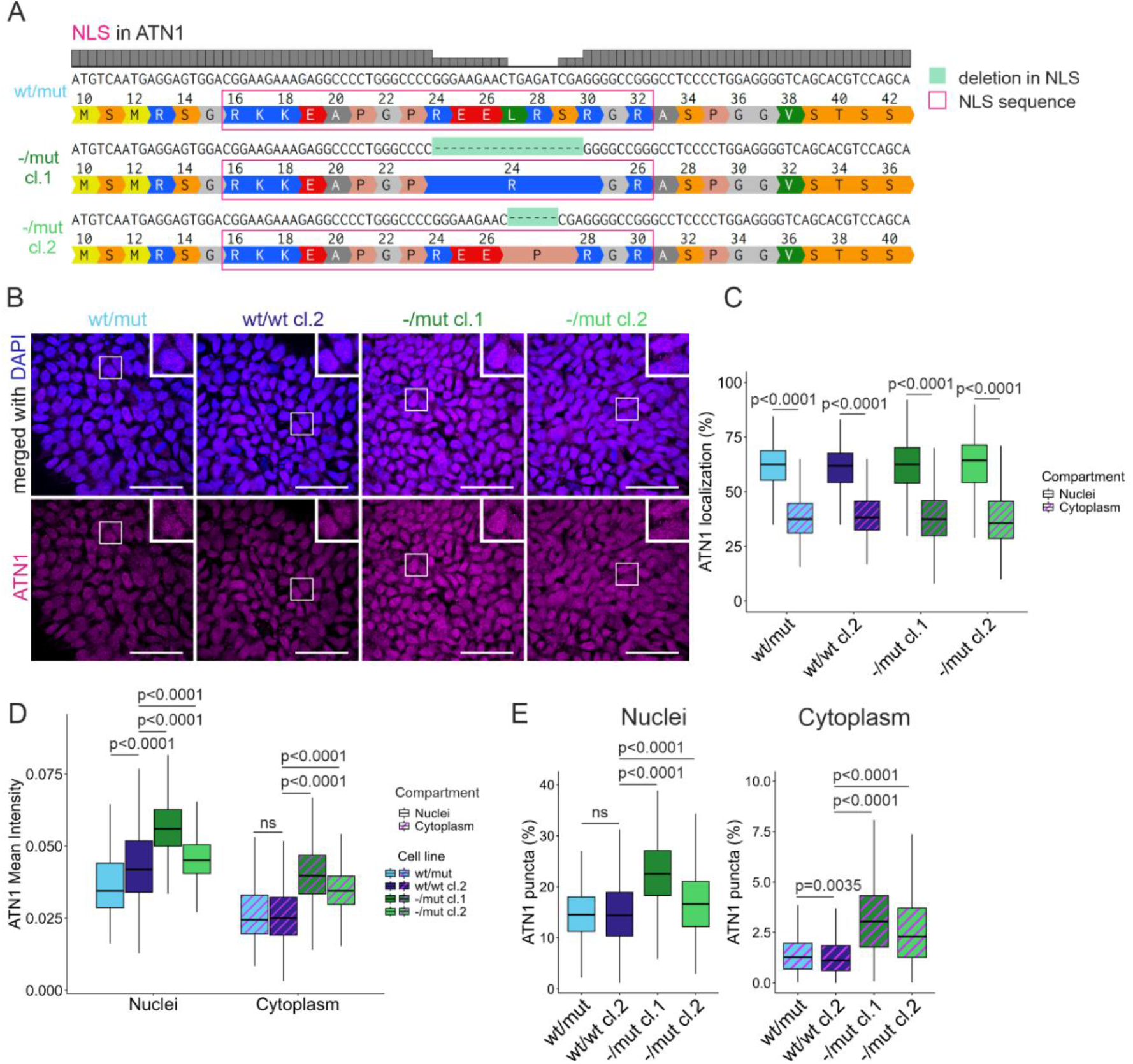
ATN1 cellular localization and puncta formation in generated iPSCs. **A**. Sequence alignment of the nuclear localization signal (NLS) region in the ATN1 comparing wt/mut and -/mut lines. The pink box indicates a canonical NLS sequence, green highlights show in-frame deletions identified in the mutated alleles of the generated -/mut cl.1 and -/mut cl.2 lines. **B**. Representative confocal immunofluorescence images showing the subcellular distribution of ATN1 (magenta) in the parental iPSC line (wt/mut), isogenic control (wt/wt cl.2), and both -/mut clones (-/mut cl.1 and -/mut cl.2). Nuclei are marked with DAPI (blue). Scale bar = 100 µm. **C**. Quantitative single-cell analysis of ATN1 mean fluorescence intensity, comparing nuclear (solid fill) and cytoplasmic (patterned fill) compartments within each analyzed iPSC line. *n* = 3 **D**. Comparison of ATN1 mean fluorescence intensity within the nuclei (left) and cytoplasm (right) across the different genotypes. *n* = 3 **E**. Quantification of the proportion (%) of ATN1 puncta localized in the nuclear (left) and cytoplasmic (right) compartments across the analyzed iPSC lines. *n* = 3 For all box plots (C–E), the horizontal line represents the median, and the box spans the interquartile range (IQR). For statistical analyses, the Kruskal-Wallis test with Benjamin-Hochberg correction was performed, and *p*-values<0.05 were considered statistically significant; *ns* – not significant. wt/mut – light blue; wt/wt cl.2 – dark blue; –/mut cl.1 –– dark green; –/mut cl.2 – light green.

Together, these data demonstrate that the presence of the wild-type allele may be crucial for maintaining the proper ATN1 pattern, whereas its absence, combined with polyQ expansion, may trigger the profound generation of granular, spot-like puncta of ATN1 protein in both cellular compartments.

### *ATN1* mutation or knock-out impairs iPSC proliferation without affecting pluripotency

Subsequently, we investigated whether ATN1 alterations affect core stemness features: cell pluripotency and self-renewal. First, we examined the synthesis of pluripotency markers across iPSC lines with various *ATN1* variants. Based on immunofluorescence staining for OCT3/4, NANOG, and SSEA4, we noted no significant difference in pluripotency markers at the protein level (**Fig. 3A, B**).

**Figure 3.**
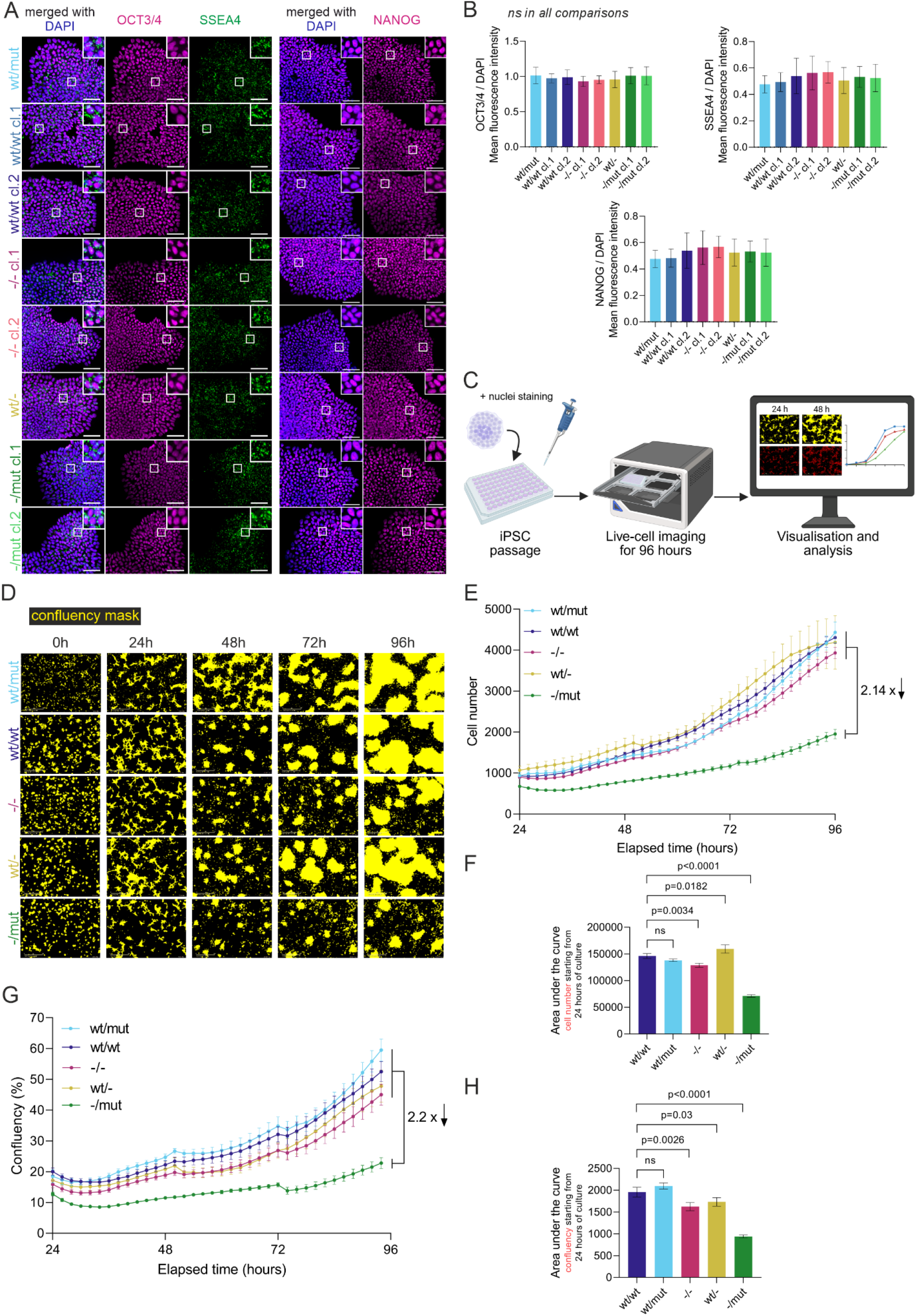
The impact of *ATN1* editing on the proliferation and pluripotency features in iPSC lines. **A**. Representative immunofluorescence images showing detection of pluripotency-associated proteins: OCT3/4 (magenta) and SSEA4 (green) in the left panel, and NANOG (magenta) in the right panel, in the parental wt/mut line and all established iPSC lines. Nuclei are marked with DAPI (blue). Scale bar = 100 µm. **B**. Quantification of immunofluorescence images with pluripotency markers (OCT3/4, SSEA4, and NANOG) in established iPSC lines. *n* = 2 **C**. Schematic workflow of the live-cell imaging assay using IncuCyte Live-Cell Analysis System. **D**.Representative images showing live-cell dynamics in a 96-hour culture of analyzed iPSCs, starting from cell seeding at 0h. The progressive cell confluency is marked by a mask (yellow). **E**. Total cell number dynamics from 24 to 94h timepoints of *in vitro* culture across all analyzed iPSC lines. An approximately 2.14-fold decrease in cell number was observed in -/mut cells compared to the mean cell number of other iPSC lines at the last timepoint (96h). *n* = 3 **F**. The iPSC growth analysis as a quantification of the area under the curve for the cell number plots obtained from live-cell imaging, calculated from 24 to 96 hours of culture. *n* = 3 **G**. Kinetic curves demonstrating a cell confluency (%) from 24 to 96h timepoints of *in vitro* culture. At the last time point (96h), the confluence of the -/mut cells decreased to approximately 2.2-fold lower than the average confluence of the other iPSC lines. *n* = 3 **H**. Quantification of iPSC confluency was shown as the area under the curve for the confluency plots obtained from live-cell imaging between 24 and 96 hours of culture. *n* = 3 For all bar plots (B, F, H), data are presented as mean ± SD. For statistical analyses, one-way ANOVA for multiple comparisons (Tukey’s test) was used, and *p*-values < 0.05 were considered statistically significant; *ns* – not significant. For all curve plots (E, G), data are presented as mean ± SEM. The dynamics of confluency and cell number over time were quantified as the area under the curve, and for statistical analyses, one-way ANOVA for multiple comparisons (Tukey’s test) was used, and *p*-values < 0.05 were considered statistically significant; *ns* – not significant. wt/mut – light blue; wt/wt cl.1 – medium blue; wt/wt cl.2 – dark blue; -/- cl.1 – dark pink; -/- cl.2 – light pink; wt/- – mustard; -/mut cl.1 – dark green; -/mut cl.2 – light green. Panels E and G show the mean values for clones representing each genotype: wt/wt – dark blue; wt/mut – light blue; -/- – dark pink; wt/- – mustard; -/mut – dark green.

Next, we evaluated the self-renewal capacity and proliferation dynamics of all iPSC lines over a 96-hour live-cell imaging culture (**Fig. 3C, D, and Supplementary Fig. 3A**). Based on confluency measurements and single-cell nuclear tracking, no differences in growth patterns were observed among the analyzed iPSC lines during the initial 24 hours post-seeding (**Supplementary Fig. 3B**). However, in subsequent times of culture, profound proliferation deficits emerged. While wt/wt and wt/mut lines maintained robust growth, one wild-type allele-expressing iPSC line (wt/-) and the *ATN1* KO (-/-) lines exhibited moderately increased and reduced proliferation rates, respectively, as compared to wt/wt iPSCs (**Fig. 3E, F, and Supplementary Fig. 3C**). The growth impairment was more profound in clones expressing only the mutated allele (-/mut) in contrast to wt/wt lines (**Fig. 3F and H**). This phenotype progressively increased over time, resulting in an approximately two-fold reduction in final cell number (2.14×) and confluency (2.2×) for the -/mut line, compared with the mean value of all other lines, by the end of the culture period (**Fig. 3E-H, and Supplementary Fig. 3C**).

Altogether, these results indicate that both the complete loss of *ATN1* and one-allele expression (-/-, wt/-) partially compromise iPSC proliferation. However, the presence of ATN1 with an expanded polyQ tract, together with the absence of a wild-type allele (-/mut), results in the most pronounced growth defect, suggesting that this severe phenotype may be caused by a combination of loss-of-function and toxic gain-of-function mechanisms.

## DISCUSSION

### Challenges in genome editing and advantages of the obtained set of lines

The present study provides a genetically defined human cellular platform for investigating the molecular functions of atrophin-1 and the consequences of distinct *ATN1* sequence alterations, including pathogenic mutation. This presents an opportunity to study early disease mechanisms within a patient-specific genetic context ^35^. By genome editing of a previously characterized DRPLA patient-derived iPSC line ^30^, we generated an allelic series comprising genetically corrected wild-type cells, complete *ATN1* knock-out cells, and heterozygous lines expressing either the wild-type or mutant allele. The most challenging part was correcting the mutant allele. Finally, two strategies were employed, resulting in two clones (**Fig. 1B**), in which DNA repair contracted the CAG tract to a length within the range of normal *ATN1* alleles in the population. Generally, isogenic correction reduces the influence of genetic background differences that complicate comparisons between unrelated patient and control iPSC lines. Such approaches are increasingly acknowledged as important for mechanistic disease modeling, including polyQ diseases ^36–38^, although clonal variation and adverse effects associated with genome editing remain important considerations ^39^. In this study, the use of two independent clones for each genotype: wt/wt, *-/-*, and -/mut further strengthens the conclusion that the observed phenotypes are related to specific *ATN1* genotype rather than a single clone-specific event.

### Expression of mutant *ATN1* without the wild-type allele causes the most prominent alterations

None of the investigated *ATN1* sequence variants substantially altered the core pluripotency state of iPSCs (**Fig. 3A, B**). These findings indicate that ATN1 is not essential for maintaining the basic molecular features of pluripotency under the culture conditions used in this study. On the other hand, all variants with at least one allele showed decreased cell number and confluence, as compared to wt/wt clone (**Fig. 3D-H**). The strongest cellular phenotypes were observed when the expanded *ATN1* allele was expressed in iPSCs, in the absence of a functional wild-type allele (-/mut lines). It is an intriguing observation that expression of only the expanded *ATN1* allele produced a more pronounced proliferation defect than *ATN1* knock-out cells. The phenotype became progressively stronger over time, with approximately two-fold reductions in final cell number and confluency in the -/mut cells (**Fig. 3D-H**). This finding suggests that the mutant protein may possess biological activity detrimental to cellular homeostasis, and in iPSCs, the absence of wild-type ATN1 unmasked this phenotype.

Cells expressing only the expanded allele showed increased ATN1 immunofluorescence signal in both the nucleus and cytoplasm, accompanied by a marked increase in ATN1 puncta (**Fig. 2D and E**). This phenotype is particularly interesting because it was detected at the pluripotency stage, and before neuronal differentiation, indicating that abnormal ATN1 protein behavior is not necessarily dependent on neuronal maturation. Further experiments will be necessary to determine the molecular nature of these punctate signals, which may represent multiple types of protein assemblies.

The altered distribution of ATN1 may reflect several, potentially overlapping mechanisms. Atrophin-1 contains regions implicated in nuclear import and export, and previous studies have shown that alterations affecting these sequences can shift the balance between nuclear and cytoplasmic localization. In particular, mutant or truncated atrophin-1 species with expanded polyQ tracts can display enhanced nuclear accumulation and cellular toxicity ^20,40^. In the present study, the -/mut clones also contained alterations that affected the predicted NLS of the mutant protein. Therefore, the increased cytoplasmic and nuclear puncta cannot be attributed exclusively to CAG expansion. It cannot be excluded that the combination of polyQ expansion and the truncated localization region may modify protein trafficking, folding, interactions with nuclear partners, or intracellular stability. Nevertheless, this study demonstrated that endogenous expression of a specific *ATN1* variant can affect iPSC proliferation.

TFs and their cofactors function as interconnected regulatory networks in which relatively modest changes in individual regulatory proteins can produce extensive downstream effects on cell state ^1,2^. Previous studies demonstrated that atrophin-1 can interact with transcriptional repression complexes ^10,41^. For example, NR2E1/TLX, a known ATN1-interacting nuclear receptor and TF, regulates neural progenitor proliferation and cell-cycle-associated genes ^10,42^. Although the present study does not establish that the proliferation phenotype is mediated directly through NR2E1/TLX, the relationship provides a plausible mechanistic framework for future investigation.

### The data support a combined effect of loss-of-function and toxic gain-of-function for ATN1 in iPSCs

Complete *ATN1* loss impairs proliferation, demonstrating that normal ATN1 function is required for optimal cellular homeostasis. At the same time, expression of the expanded allele in the absence of wild-type ATN1 produces a substantially stronger phenotype than ATN1 loss alone. This argues that in iPSCs, the mutant protein is not simply an inactive allele but retains properties that perturb cellular function. The data demonstrate that the presence of a functional *ATN1* allele substantially influences the behavior of the mutant protein. The relatively less pronounced phenotype in wt/mut cells suggests that wild-type atrophin-1 may buffer the effects of the expanded protein, either by maintaining appropriate transcriptional complexes, facilitating protein turnover, or limiting aberrant protein interactions.

Interestingly, we and others previously investigated human stem cells at early stages of neural differentiation in the context of the huntingtin (*HTT*) gene, and found that selected phenotypes were common for HD and *HTT* knock-out cells ^43–45^. The present results, although based solely on iPSC phenotypes, support the assumption that polyQ diseases may have a loss-of-function component and show alterations at the early stages of cell development ^46–48^. Moreover, clinical distinction between DRPLA and CHEDDA suggests that different structural alterations of ATN1 can perturb distinct molecular functions, connected with abnormal processing and altered protein interactions or regulatory activity. The present allelic series provides an experimental framework for understanding ATN1 function and dysfunction.

### Limitations of the study and future perspectives

The experiments were performed in iPSCs, which provide an experimentally relevant human model but do not allow for direct reference to effects in differentiated cells. As DRPLA is primarily a neurodegenerative disorder, some disease mechanisms may only emerge after neuronal induction. Therefore, the next step should consequently be direct induction of the obtained set of lines into relevant neuronal populations. Comparing, e.g., neuronal differentiation efficiency and neuronal subtype specification between generated lines would directly test whether the molecular alterations detected in iPSCs translate into neuronal phenotypes. Obtaining and integrating transcriptomic data with ATN1 chromatin occupancy and proteomic interactome analyses should also help to describe ATN1’s direct regulatory network.

In conclusion, this study establishes a comprehensive set of human iPSC lines to dissect ATN1 biology and demonstrates that ATN1 functions prior to cell differentiation. The preservation of pluripotency, despite pronounced changes in proliferation in *ATN1* mutant lines, indicates that ATN1 is not a core pluripotency factor but instead contributes to the regulatory processes that maintain appropriate cellular growth and developmental competence. The marked phenotype of cells expressing only mutant *ATN1* further supports a combined loss-of-function and toxic gain-of-function mechanism. Thus, the generated isogenic iPSC panel represents a valuable foundation for future studies aimed at defining ATN1-dependent regulatory networks, understanding the molecular basis of both DRPLA and CHEDDA, and identifying disease-relevant pathways during neuronal differentiation that could contribute to neurodegeneration. Despite advances in preclinical testing of disease-modifying treatments for DRPLA ^49,50^, bridging basic biological insights with clinical data is essential to fully understand disease complexity ^51,52^.

## Acknowledgments

We acknowledge the use of the infrastructure developed under the project NEBI - National Research Center for Imaging in the Biological and Biomedical Sciences, POIR.04.02.00-00-C004/19, co-financed through the European Regional Development Fund (ERDF) in the frame of Smart Growth Operational Programme 2014-2020 (Measure 4.2 Development of modern research infrastructure of the science sector). We acknowledge the Laboratory of Single Cell Analysis and Laboratory of Cell and Tissue Culture, Laboratory of Mammalian Model Organisms ICHB PAS for access to equipment. We would like to thank the lab members for the discussions and their support of this study. Figures were created using biorender.com and GraphPad Prism.

## Author Contribution Statement

Conceptualization: AF, EK, NZ, BN. Experiments and data analysis: EK and NZ with contributions from BN, PJ, MDM, MS, OB, AF. Genome editing strategy: MDM, MO. Funding acquisition: AF, BN. Project supervision: AF. Manuscript draft and figures: NZ with contributions from AF and EK. All authors reviewed and approved the final manuscript.

