## Supplementary Files for "Modeling *ATN1* mutations in human iPSCs reveals early developmental roles of atrophin-1"

### SUPPLEMENTARY FIGURES

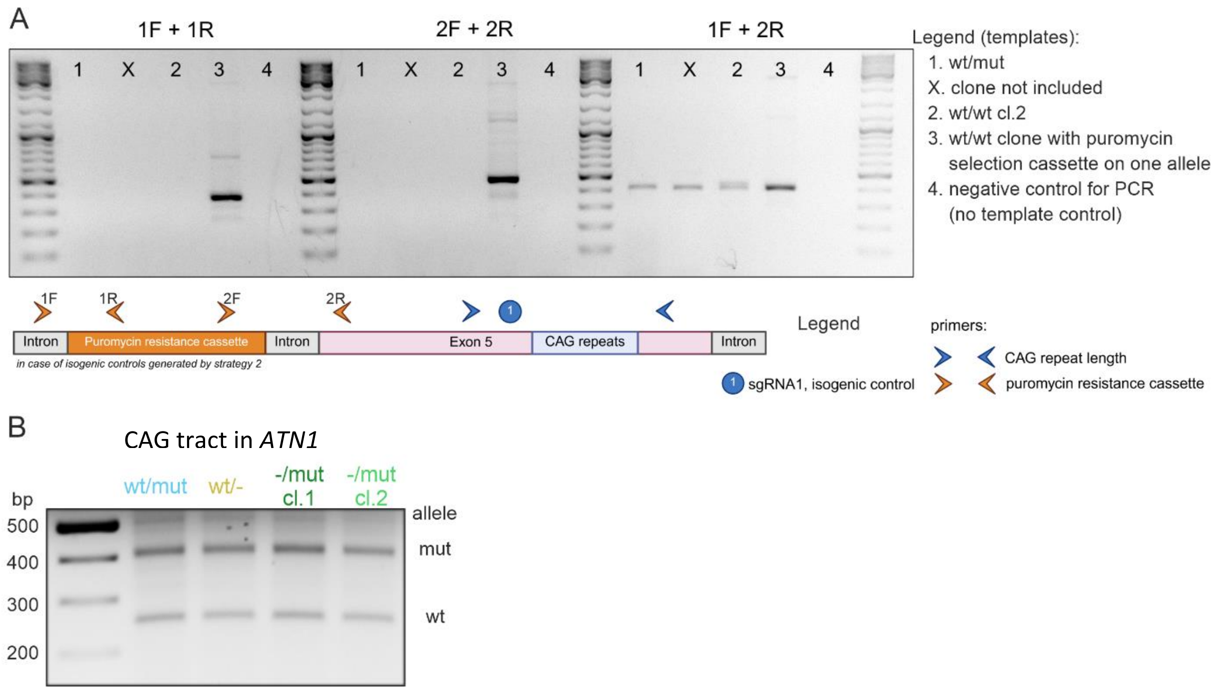


**Supplementary Figure 1. Validation of selected sequences in the *ATN1* locus in the generated iPSC lines**

- **A.** Genotyping PCR analysis testing the occurrence of the puromycin resistance cassette in the targeted *ATN1* intronic region. The upper panel shows the gel electrophoresis of PCR products amplified with specific primer combinations (1F+1R, 2F+2R, 1F+2R) for parental wt/mut (1), generated control wt/wt cl.2 (2), intermediate puromycin-resistant wt/wt clone (3), and negative control (4) templates. This result confirmed the removal of the puromycin resistance cassette in wt/wt cl.2. The schematic below illustrates the *ATN1* locus, including the puromycin cassette and primer binding sites.
- **B.** PCR amplification of the genomic *ATN1* fragment spanning the CAG repeat tract in wt/mut and single allele knock-out iPSCs (wt/-, -/mut cl.1, and -/mut cl.2). The gel confirms the presence of the retained wild-type (wt) and expanded mutant (mut) allele at the genomic level.

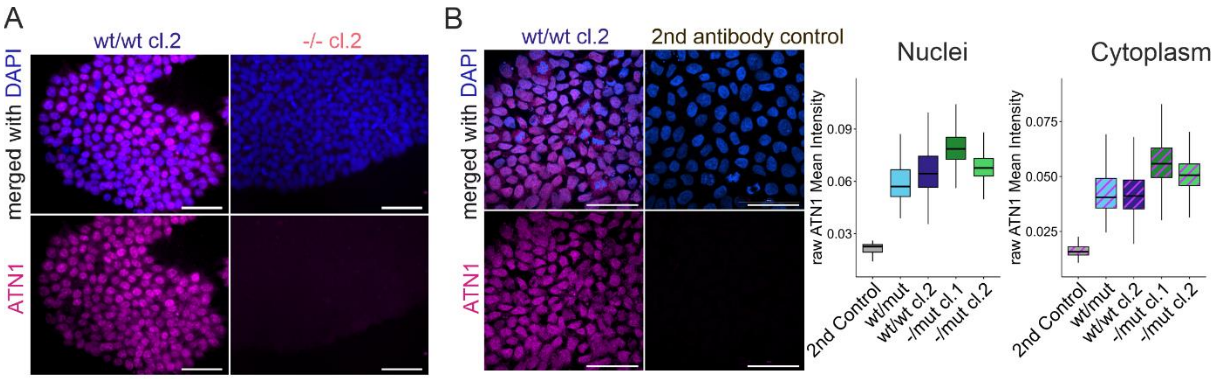


**Supplementary Figure 2. Controls for ATN1 immunofluorescence staining**

- **A.** Representative images for ATN1 protein detection in wt/wt cl.2 and -/- cl.2 iPSC lines. Scale bar = 100 µm.
- **B.** Representative images with ATN1 immunostaining of wt/wt cl.2, and only secondary antibody as a control for non-specific binding and background fluorescence. Quantitative single-cell analysis of ATN1 mean fluorescence intensity, showing background fluorescence signal from secondary antibody control (Control, gray) compared to analyzed iPSC lines in nuclear (left) and cytoplasmic (right) compartments. wt/mut – light blue; wt/wt cl2. – dark blue; -/mut cl1. – dark green; -/mut cl.2 – light green. The horizontal line represents the median, and the box spans the interquartile range (IQR).

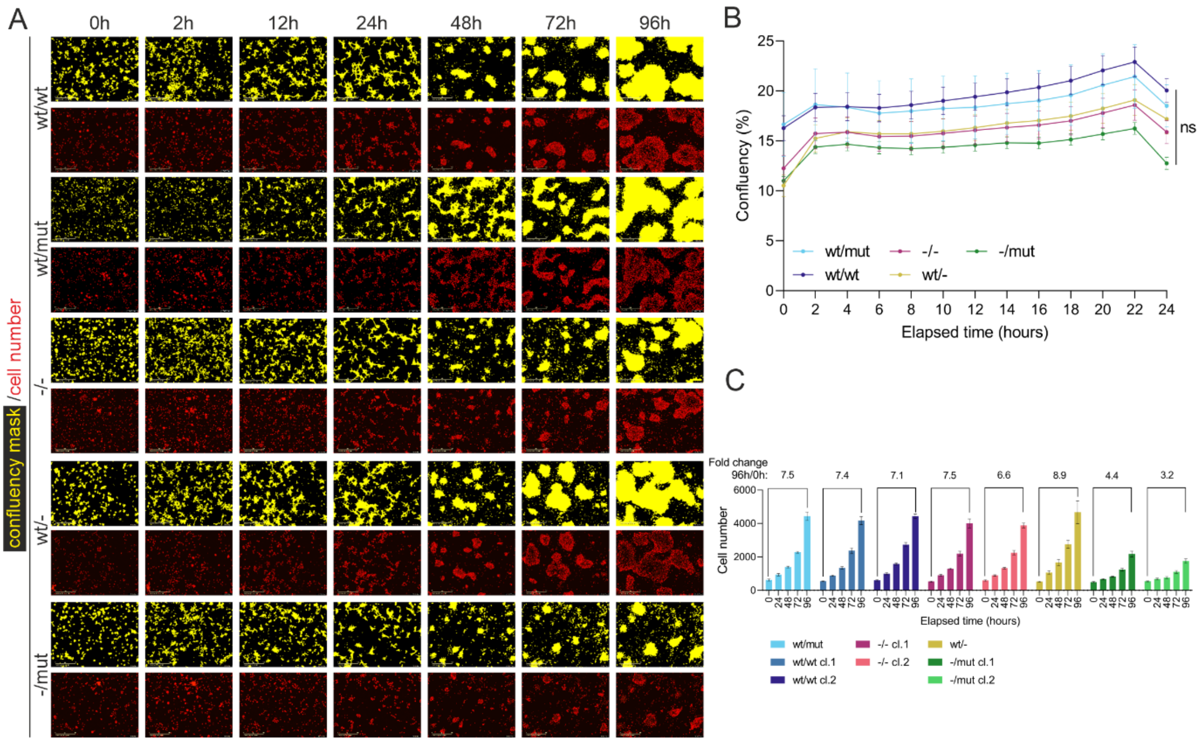


**Supplementary Figure 3. Additional results for live-cell analysis of cell confluency and proliferation dynamics in *ATN1*-edited iPSCs**

- **A.** Representative images showing live-cell analysis in a 96-hour *in vitro* culture of the generated iPSC ATN1 panel, timepoints: 0h, 2h, 12h, 24h, 48h, 72h, 96h. The cell confluency (%) is marked by a yellow mask (upper rows), and the cell number is marked by a red mask (lower rows).
- **B.** Analysis of confluency (%) during the initial 24 hours of culture. The data are presented as mean values ± SEM. Plotted genotypes: wt/wt – dark blue; wt/mut – light blue; -/- – dark pink; wt/- – mustard; -/mut – dark green. For statistical analysis, the last time point (24h) was considered, and a one-way ANOVA with multiple comparisons (Tukey's test) was performed. The $p$-values < 0.05 were considered statistically significant; *ns* – not significant. $n = 3$
- **C.** Bar graphs showing the cell number progression at 24-hour intervals (from 0 to 96 hours) for each analyzed iPSC line. The numeric values above the brackets indicate the calculated fold change in cell number between 96h and 0h. Data are presented as mean ± SD. For statistical analysis, the last time point (24h) was considered, and a one-way ANOVA with multiple comparisons (Tukey's test) was performed. The $p$-values < 0.05 were considered statistically significant; *ns* – not significant. $n = 3$. wt/mut – light blue; wt/wt cl.1 – medium blue; wt/wt cl.2 – dark blue; -/- cl.1 – dark pink; -/- cl.2 – light pink; wt/- – mustard; -/mut cl.1 – dark green; -/mut cl.2 – light green.

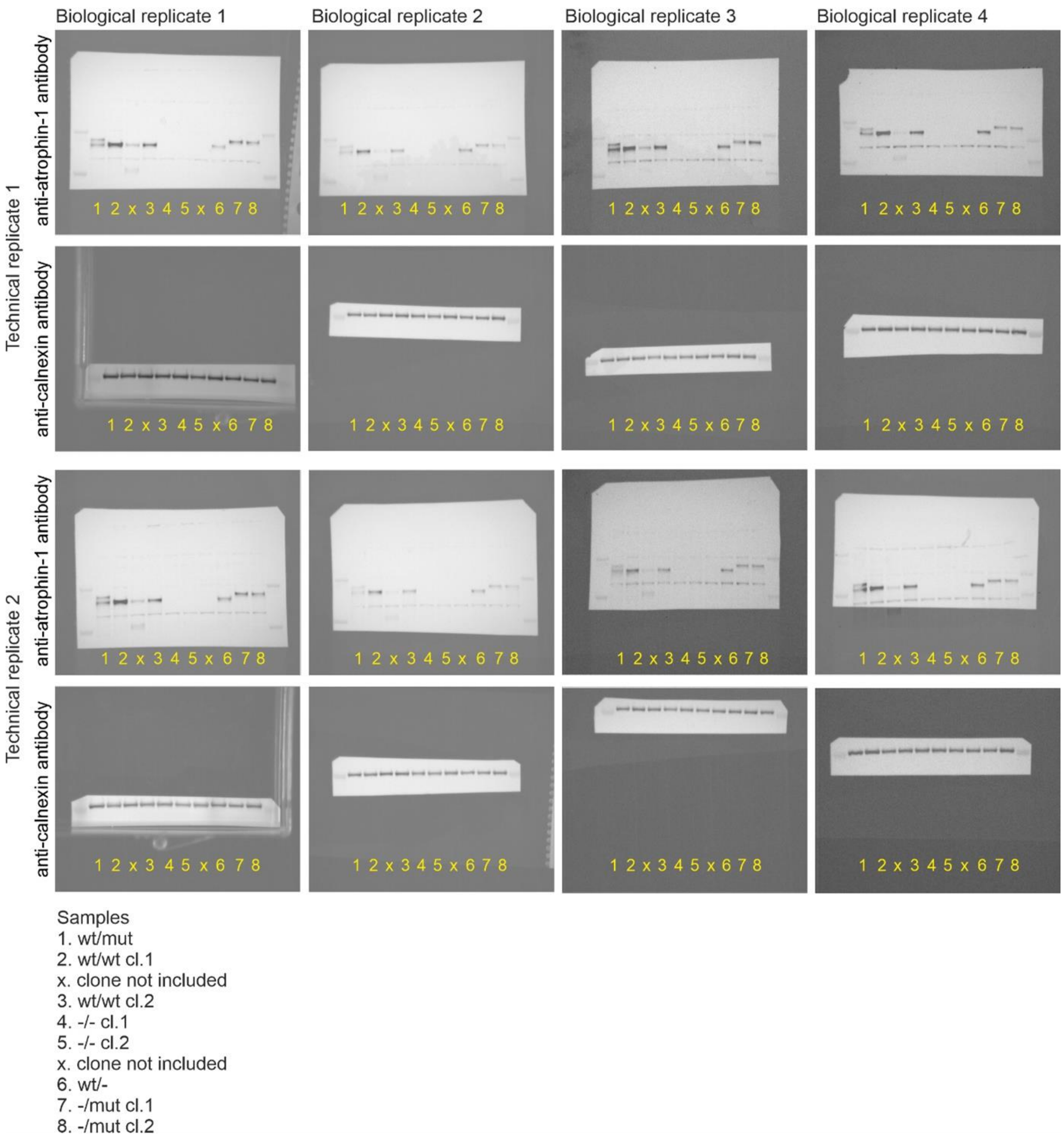


**Supplementary Figure 4. Full membranes for western blot analysis of ATN1 protein level in iPSC lines**

Atrophin-1 and calnexin (loading control) were detected in four biological replicates, each comprising two technical replicates. Sample identities are indicated below the blots. The blots were used for ATN1 level quantification presented in Fig.1 in panels E, H, and J. Selected blots were used for presentation of exemplary results shown in Fig.1 D, G, and I.

### SUPPLEMENTARY MATERIALS AND METHODS

#### Genome editing and clonal selection

*Strategy 1: RNP complex and HDR*

*Generated iPSC lines: wt/wt cl.1*

The ribonucleoprotein (RNP) complex was assembled following the manufacturer's instructions for the IDT Alt-R CRISPR-Cas9 System. Briefly, crRNA containing the target sequence and tracrRNA were mixed at equimolar concentrations to obtain a final duplex concentration of 60 µM, heated at 95 °C for 5 min, and then incubated at room temperature for 20 min. Subsequently, 60 pmol of the guide RNA duplex was combined with 30 pmol of SpCas9 protein (VBCF Protein Technologies Facility, Austria) and incubated for 20 min at room temperature to form the RNP complex. The assembled RNP and donor template (Supplementary Table 1) were delivered into patient-derived DRPLA iPSCs (wt/mut) by electroporation using the Neon Transfection System (Invitrogen, USA). Upon reaching approximately 70% confluency, iPSCs were dissociated into single cells using Accutase solution (Sigma-Aldrich, USA), and $2 \times 10^5$ cells were electroporated with the RNP complex using the following parameters: 1100 V, 20 ms, 2 pulses. Following electroporation, cells were plated onto Geltrex-coated wells of a 6-well plate in StemFlex medium (Gibco, USA) supplemented with 10 µM Y-27632 (StemCell Technologies, Canada) and cultured for the next 2 days. The cells were then replated at single-cell density on Geltrex-coated 6-mm dishes and cultured for an additional 7 days. Individual colonies were manually picked and expanded in 96-well plates. To identify clones, genomic DNA was extracted from each monoclonal line using QuickExtract DNA Extraction Solution (Lucigen, Middleton, WI, USA), and PCR amplification was performed with primers flanking the CRISPR-edited region, followed by gel electrophoresis and Sanger sequencing.

The primary objective of strategy 1 was to correct the mutation through homology-directed repair (HDR) using a donor template. Unexpectedly, the resulting clone exhibited a contraction of the CAG repeat tract without the use of the donor template. Since the length of the corrected CAG repeat tract matched that of the wild-type allele, we hypothesize that the wild-type allele served as the repair template for the mutant allele.

*Strategy 2 and 3: pSpCas9-2A-GFP plasmid with or without a piggyBac-based approach*

*Generated iPSC lines: wt/wt cl.2, -/- cl.1, -/- cl.2, wt/-, -/mut cl.1, -/mut cl.2*

To facilitate the generation of isogenic controls, we employed a CRISPR-Cas9 and *piggyBac* transposon-based approach to promote HDR. In this approach, 1.784-kbp and 2.971-kbp homologous arms derived from the *ATN1* gene (coordinates for homologous arms GRCh38: chr12:6,933,505-6,938,239) were cloned into the pJOP plasmid (Addgene plasmid #87228) flanking the *piggyBac* transposon cassette containing GFP and puromycin-resistance gene, using the NheI/NotI and BsrGI/SpeI restriction sites, respectively. The 2.971-kbp homologous arm contained a 15 CAG repeat tract. The BsrGI restriction site was introduced into the pJOP plasmid immediately downstream of the *piggyBac* left (5') inverted repeat by site-directed mutagenesis according to the manufacturer′s instructions (QuickChange II Site-Directed Mutagenesis Kit, Agilent). The resulting plasmid was designated pJOP-ATN1-CAG and used for subsequent experiments. The sgRNA1 targeting exon 5 of the *ATN1* gene, required for DSB induction and initiation of HDR, and sgRNA2, targeting exon 3 of the *ATN1* gene for knock-out generation, were designed using CRISPOR software (http://crispor.tefor.net/crispor.py). Next, they were cloned into the pSpCas9-2A-GFP plasmid (Addgene plasmid #48138). Briefly, the top and bottom strands of 20-nt guide RNA were synthesized (IBB, Warsaw), annealed and ligated into the BpiII-digested pSpCas9(BB)-2A-GFP (PX458) plasmid. (Addgene, Cambridge, MA, USA). Ligated products were transformed into chemically competent E. coli GT116 cells (InvivoGen, San Diego, CA, USA), and the cells were plated onto ampicillin selection plates (100 µg/mL ampicillin) and incubated at 37°C overnight. Plasmid DNA was isolated using the Gene JET Plasmid Miniprep kit (Thermo Scientific) and verified with Sanger sequencing. For larger-scale plasmid preparations, the Qiagen Midi kit was used (Qiagen, Hilden, Germany). The *ATN1* sequences complementary to each sgRNA are presented in Supplementary Table 1.

The pJOP-ATN1-CAG together with pSpCas9-2A-GFP containing sgRNA1 or pSpCas9-2A-GFP with sgRNA2 were delivered into DRPLA iPSCs (wt/mut) via electroporation using the Neon Transfection System (Invitrogen, USA). At approximately 70% confluency, cells were dissociated into single-cell suspensions with Accutase (Sigma-Aldrich, USA), and $1 \times 10^6$ cells were electroporated with 4 µg of each plasmid using the following parameters: 1400 V, 20 ms, 1 pulse. After electroporation, cells were plated on Geltrex-coated wells of a 6-well plate in StemFlex medium (Gibco, USA) supplemented with 10 µM Y-27632 (StemCell Technologies, Canada) and cultured for the next 2 days. Subsequently, puromycin selection or fluorescence-activated cell sorting (FACS) was applied to enrich the CRISPR-edited pool of isogenic control or knock-out clones, respectively. After 3 days of culture, cells were replated at single-cell density on Geltrex-coated 6-mm dishes and maintained for an additional 7 days. Individual colonies were manually picked and expanded in 96-well plates. Genomic DNA was isolated from each monoclonal line using the Genomic DNA Isolation Kit (Norgen Biotek, Canada), and PCR amplification of the region flanking the CRISPR-edited site, followed by agarose gel electrophoresis and Sanger sequencing, was performed to confirm the genotype of each clone. To remove the selection cassette from the genome of the isogenic control $1.5 \times 10^6$ iPSC line, 2 µg of *piggyBac* excision-only vector (System Biosciences, USA) was used, followed by negative selection at 0.25 µM fialuridine and clonal selection. Moreover, during the generation of *ATN1* knock-out iPSC lines, one clone carrying a knock-out of the mutant allele (containing 68 CAG repeats) and two clones with a knock-out of the wild-type allele (containing 16 CAG repeats) were identified.

### SUPPLEMENTARY TABLES

**Supplementary Table 1**. Target sequences for sgRNAs (complementary *ATN1* sequence) and HDR donor template for strategy 1

| Name | Sequence (5'-3') | Target | Application |
|---|---|---|---|
| sgRNA1 | TGATGGTGATGTGTTGAGAC | *ATN1* exon 5 | Isogenic control |
| sgRNA2 | CCCGGGAAGAACTGAGATCG | *ATN1* exon 3 | *ATN1* knock-out |
| HDR donor template | AACAGCAATGCCCATCCAGG CCCCTTCCCTCCCTCTACTGG GGCCCAGTCCACCGCCCACCC ACCAGTTTCAACACATCACCA TCACCACCAGCAACAGCAAC AGCAGCAGCAGCAGCAGCAG CAGCAGCAGCAGCAGCAGCA GCAGCATCACGGAAACTCTG GGCCCCCTCCTCCTGGAGCAT TTCCCCACCCACTGGA | *ATN1* exon 5 | Isogenic control |

**Supplementary Table 2.** List of antibodies used for immunostaining

| Antibodies | Catalog number | Vendor | Species | Dilution |
|---|---|---|---|---|
| anti-NANOG | MA1-017 | Thermo Fisher Scientific | mouse | 1:100 |
| anti-SSEA4 | 4755 | Cell Signaling | mouse | 1:100 |
| anti-OCT3/4 | PA5-27438 | Thermo Fisher Scientific | rabbit | 1:200 |
| anti-ATN1 | HPA031619 | Sigma Aldrich | rabbit | 1:150 |
| anti-mouse, AlexaFluor488-conjugated | 715-545-151 | Jackson ImmunoResearch | donkey | 1:400 |
| anti-mouse, TRITC-conjugated | 715-025-150 | Jackson ImmunoResearch | donkey | 1:400 |
| anti-rabbit, TRITC-conjugated | 711-025-152 | Jackson ImmunoResearch | donkey | 1:400 |

**Supplementary Table 3.** List of antibodies used for western blot

| **Antibodies** | **Catalog number** | **Vendor** | **Species** | **Dilution** |
|---|---|---|---|---|
| anti-ATN1 | HPA031619 | Sigma Aldrich | rabbit | 1:1000 |
| anti-CANX | C4731 | Merck | mouse | 1:10000 |
| anti-mouse, peroxidase-conjugated | 715-035-150 | Jackson ImmunoResearch | donkey | 1:2000 |
| anti-rabbit, peroxidase-conjugated | 711-035-152 | Jackson ImmunoResearch | donkey | 1:2000 |

**Supplementary Table 4.** List of primers used in the study

| **Name** | **Sequence (5'-3')** | **Target** |
|---|---|---|
| *ATN1*_CAG_F | CTTAGCCAACAGCAATGCCC | *ATN1* exon 5 |
| *ATN1*_CAG_R | AGACATGGCGTAAGGGTGTG | *ATN1* exon 5 |
| 1_F | AAACCAAGGGCTTGGGGGAT | *ATN1* intron sequence, upstream of the puromycin-GFP cassette |
| 1_R | CGTAATTACATCCCTGGGGGC | 5' region of the puromycin-GFP cassette |
| 2_F | CATTGACAAGCACGCCTCAC | 3' region of the puromycin-GFP cassette |
| 2_R | TCATCATTAAGGCTCCGCCC | *ATN1* exon 5 sequence, downstream of the puromycin-GFP cassette |
| *ATN1*_KO_F | AGACATGAAAGATGAGGGGCG | *ATN1* exon 3 |
| *ATN1*_KO_R | CACCGTTACCCTCTGCCAAA | *ATN1* exon 3 |
